# A novel full-reduced HMGB1 and kartogenin-containing bio-active scaffold for meniscus regeneration

**DOI:** 10.1101/2020.03.26.010108

**Authors:** Hongyao Xu, Zhihong Dai, Xiangjie Zou, Pengcheng Xia, Mohammad Ahmad Aboudi, Jingwen Wang, Warren Alexander Lam Sung Foon, He Huang

**Affiliations:** Department of Orthopaedics, Nanjing First Hospital, Nanjing Medical University, 68 Changle Road, Nanjing, Jiangsu 210006, China; China Orthopaedic Regeneration Medicine Group, Hangzhou, Zhejiang 310000, China

**Keywords:** HMGB1, kartogenin, alginate scaffold, cell migration, meniscus regeneration

## Abstract

A novel bio-active scaffold for enhancing wounded meniscus healing has been developed by combination of High mobility group box 1 protein (HMGB1) and kartogenin (KGN) with alginate gel. The properties of the bio-active scaffold were also investigated using an *in vitro* cell culture model and *in vivo* rat wounded meniscus healing model. This HMGB1-KGN-containing bioactive scaffold released HMGB1 and KGN into wound area and kept high concentrations of HMGB1 and KGN in the system for more than two weeks. This HMGB1-KGN-containing bioactive scaffold also activated rat bone marrow stem cells (BMSCs) from G_0_ to G_Alert_ stage and promoted cell proliferation as evidenced by 5-bromo-2-deoxyuridine (BrdU) incorporation testing. Our results also demonstrated that the HMGB1-KGN-containing bioactive scaffold induced cell migration *in vitro* and recruited the cells to wound area to promote wounded rat meniscus healing *in vivo*.

## Introduction

The menisci are C-shaped fibrocartilage located between the femoral condyles and tibial plateau to transmit the load across the joint space [1]. Menisci play an important role in joint stabilization, proprioception, lubrication and protection of articular cartilage [2]. Such environment and function lead to meniscus easily injury and damaged meniscus can cause the progression of cartilage degeneration and result in osteoarthritis [3]. Clinical studies have found that it is difficult to regenerate the meniscus due to the specific structural and functional properties of the meniscus tissue [1].

Histology studies have shown that meniscus is a geometrically and biochemically complex tissue. Its outer 1/3 tissue is formed by collagen type I produced by fibroblast-like cells, while its inner 2/3 tissue is formed by collagen II and proteoglycan produced by chondrocyte-like cells [4]. Vascularization in meniscus is of high relevance. The meniscus is fully vascularized during fetal development though blood supply diminishes over time, receding to the peripheral 20-30% by 10 years of age [4], [1].

The injury in the avascular zone of the meniscus is generally more complex and broad, and is often associated with a poor prognosis following repair [5]. Promoting the healing process in the avascular zone of the meniscus is an ongoing challenge for both clinical medical doctors and orthopedic researchers [6]. Recently, tissue engineering approaches that involve the use of adult tissue-derived multi-potent stem cells and biomaterial scaffolds have gained increasing attention for enhancing the wounded meniscus healing. However, the isolation and culture of stem cells not only need the donor tissue obtained by additional surgical procedure but also take long time and increase contamination risk. Therefore, recruiting stem cells to wound area to enhance wound healing is still a new challenge for orthopedic researchers.

High mobility group box 1 protein (HMGB1) is a ubiquitously expressed, highly conserved nuclear protein [7]. In normal circumstances, HMGB1 is almost always present in the nuclei of mammalian cells and plays an important role in various biological processes including transcription, DNA repair, cell differentiation and development [8]. When the cells are stimulated or damaged, HMGB1 can be released from the nucleus to the cytoplasm [9]. The extracellular HMGB1 can interact with some cellular receptors and surface molecules to regulate the cell proliferation and cell migration [7]. It has been found that HMGB1 has two different activities due to its two redox sensitive cysteine moieties in its 215 amino acid structure [10]. When these cysteine residues in HMGB1 are in reduced thiol form, this kind of HMGB1 is called as fully reduced HMGB1 (frHMGB1) which is a chemoattractant of motile cells. However, when these thiols are oxidized to form a disulfide bond which is called oxidized HMGB1 (oxHMGB1). The oxHMGB1 is an inducer of cytokines. It has been reported that frHMGB1 binds to chemokine (C-X-C motif) ligand 12 (CXCL12) to form a HMGB1-CXCL12 heterocomplex which interacts with CXCR4 to promote the migration of monocytes and fibroblasts [11]. The recent study has shown that frHMGB1 can transitions stem cells from G_0_ to G_Alert_ [12].

Another key point for meniscus regeneration is to induce chondrogenesis differentiation of the stem cells. Recent studies have found that kartogenin (KGN), a small compound can promote the chondrogenic differentiation of human and mouse mesenchymal stem cells (MSCs) [13]. Several studies have used KGN to enhance wounded cartilage regeneration [14, 15]. We have shown that KGN-treated autologous tendon graft can enhance the wounded rat meniscus healing [16].

Finally, a suitable scaffold is needed to deliver HMGB1 and KGN to the wound area which can release appropriate concentrations of HMGB1 and KGN to enhance wound healing. It is well known that alginate is a naturally occurring anionic polymer and has been used for many biomedical applications due to its biocompatibility, low toxicity, and low cost [17].

In this study, we developed a novel bio-active scaffold by the combination of fully reduced HMGB1 (frHMGB1) and KGN with alginate hydrogel for promoting rat wounded meniscus healing. Our hypotheses are that 1) this bio-active scaffold can deliver frHMGB1 and KGN directly to the wound area and release appropriate concentrations of frHMGB1 and KGN in the wound area; 2) frHMGB1 induces stem cell migration *in vitro* and recruits the stem cells to the wound area *in vivo*; 3) The inhibition of HMGB1 activity by HMGB1 inhibitor can block the cell migration and slow down the wound healing; 4) KGN can induce the stem cells to differentiate into chondrocytes to enhance the wounded meniscus healing; 5) HMGB1 and KGN-containing alginate hydrogel is a suitable bioactive scaffold for promoting wounded meniscus regeneration.

## Materials and methods

### Animals

Twenty-seven Sprague Dawley (SD) rats (3 months old, female) and three GFP transgenic SD rats (3 months old, female) were used in this study. The experiments were done at Nanjing Medical University (NMU) following the approved protocol by the Institutional Animal Care and Use Committee (IACUC) of NMU.

### The preparation of HMGB1 and KGN-containing bioactive scaffold

Sodium alginate (Sigma, St Louise, USA) was dissolved with distilled water to make a 2% of alginate solution (AS). The frHMGB1 was dissolved with distilled water to make a 1 μg/μl of frHMGB1. KGN (Sigma, St Louise, USA) was dissolved with DMSO to make a 25 mg/ml of KGN solution. All solutions were filtered through a 0.2 μm membrane under sterile conditions. The bioactive scaffold for enhancing wounded meniscus healing was prepared by adding 10 μl of frHMGB1 solution (10 μg frHMGB1 in 10 μl water) and 10 μl of KGN solution (250 μg of KGN in 10 μl of DMSO) into 80 μl of alginate solution (AS) and mixed well to get a HMGB1-KGN-AS solution. Each 10 μl of HMGB1-KGN-AS solution was dropped into 2% of CaCl_2_ solution to obtain a HMGB1-KGN-containing bioactive scaffold which contained 1 μg of frHMGB1 and 25 μg KGN. Similarly, the control scaffold was made by adding 10 μl of distilled water and 10 μl of DMSO into 80 μl of alginate solution and mixed well to make a DMSO-AS solution. Each 10 μl of DMSO-AS solution was dropped into 2% of CaCl_2_ to make a DMSO-containing control scaffold. Furthermore, 10 μl of KGN solution and 10 μl of distilled water were added into 80 μl of AS and mixed well to make a KGN-containing scaffold by dropping 10 μl of KGN-AS mixture into 2% CaCl_2_ solution. Moreover, 10 μl of frHMGB1 solution and 10 μl of DMSO were added into 80 μl of AS and mixed well to make a HMGB1containing scaffold by dropping 10 μl of HMGB1-AS mixture into 2% CaCl_2_ solution.

### The determination of HMGB1 released from HMGB1-containing scaffold

Each HMGB1-containing scaffold prepared by above procedures was added in 1 ml of PBS and the concentration of HMGB1 in PBS was determined at different time point using an ELISA kit (Tecan Group Ltd., Mannedort, Switzerland) according to the manufacture’s protocol. Three samples were tested at each time point.

### The determination of KGN released from KGN-containing scaffold

Each KGN-containing scaffold prepared by above procedures was added in 1 ml of PBS and the concentration of KGN in PBS was determined at different time point using high performance liquid chromatography (HPLC; Hitachi, Japan) according to the published protocol [14].

### Isolation of bone-marrow derived stem cells (BMSCs)

The BMSCs were isolated from femur bones of either SD rats or GFP transgenic SD rats according to the published protocols [18]. Briefly, a needle (21-guage) fastened to a syringe containing 0.1 ml of heparin (1000 units/ml) to aspirate 1 ml of bone marrow followed by washing the aspirates twice with phosphate-buffered saline (PBS). The bone marrow-PBS solution was centrifuged at 1500 g for 5 minutes. After discarding the supernatant, the cells were re-suspended in 20% fetal bovine serum-containing Dulbecco’s Modified Eagle Medium (DMEM) with 1% of penicillin and streptomycin and incubated at 37°C with 5% of CO_2_ and 95% of air atmosphere.

### The effect of HMGB1 on inducing the migration of rat BMSCs

The effect of HMGB1 on inducing the migration of rat BMSCs were examined using a Trans-well plate (Millipo Sigma, St Louise, USA). The rat BMSCs in DMEM at passage 1 were seeded into the upper layer of insert well of the trans-well plate. The lower layer of the trans-well plate was filled with various concentrations of HMGB1 (1-100 ng/ml; Tecan Group Ltd., Mannedort, Switzerland)-containing DMEM or a HMGB1-AS scaffold-containing (1000 ng HMGB1/scaffold/ml) DMEM. After 24 hours, the cells migrated through the membrane to the lower layer well were counted by H33342 staining (Sigma, St Louise, USA).

### The inhibition effect of HMGB1 inhibitors on cell migration induced by HMGB1

The effect of HMGB1 on inducing the migration of rat BMSCs were further examined with HMGB1 inhibitors. The rat BMSCs in DMEM at passage 1 were seeded into the upper layer of insert well of the trans-well plate. The lower layer of the trans-well plate was filled with 100 ng/ml of HMGB1-containing DMEM with various concentrations of FSP-ZM1 or AMD3100 (Sigma, St Louise, USA). After 24 hours, the cells migrated through the membrane to the lower layer of each well were counted by H33342 staining (Sigma, St Louise, USA).

### The effect of HMGB1 on cell activation from G_0_ to G_Alert_

The effect of HMGB1 on cell proliferation was examined by BrdU incorporation testing. The rat BMSCs at passage 1 were seeded into 12-well plate (2 × 10^4^ cells/well) and cultured with 20% FBS-DMEM for 3 days. Then the cells were cultured in serum-free medium for another 24 hours, finally various concentrations of HMGB1 (1-100 ng/ml) and a HMGB1 (1000 ng)-containing AS scaffold with 10 μM of 5-bromo-2’-deoxyuridine (BrdU; Sigma, St Louise, USA) were added into 1 ml of serum-free medium and cultured for additional 24 hours. The cell numbers incorporated BrdU were determined by immunostaining with anti-BrdU antibody (Sigma, St Louise, USA). The positive cells were identified by Cy3-conjugated second antibody, and counted by semi-quantification.

### Semi-quantification of BrdU positive cells

The activated cells induced by HMGB1 were determined by BrdU incorporation testing. Briefly, after the treatment with HMGB1 and BrdU, the BrdU-labeled DNA was determined under a fluorescent microscope (Nikon Eclipse, TE2000-U). Four images of each well were randomly taken and the BrdU incorporated cells were stained with red fluorescence. The proportion of positive stained cells in each image was calculated by dividing the red fluorescent cell numbers by total cell numbers stained with H33342 (blue fluorescent cells). The average of four images from each well was used to represent the percentage of positive staining, which is the extent of BrdU incorporated cells in the respective media.

### In vivo model for wounded rat meniscus healing

The meniscus cells were eliminated from total 24 SD rats by the treatment of the knees with 6 Gy X-ray, then a wound was created in the meniscus with a 1 mm diameter biopsy punch. The wounded rats were divided into four groups with 6 rats in each group. The wound in group-1 was treated with a DMSO-containing scaffold (**DMSO**); the wound in group-2 was treated with a KGN-containing scaffold (**KGN**); the wound in grou-3 was treated with a HMGB1-containing scaffold (**HMGB1**); the wound in group-4 was treated with a HMGB1-KGN-containing scaffold (**HMGB1-KGN**). The GFP-labeled BMSCs (10^6^ cells) isolated from GFP rats were injected into each wounded rat via tail vein. Three rats from each group were sacrificed 2 weeks post-operation and three rats were sacrificed 4 weeks post-operation. The menisci from left knee of each rat were used for cellular analysis, and the menisci from right knee of each rat were used for histology analysis.

### GFP-labeled cell isolation from wounded rat menisci

The wound area tissues were collected from the menisci of the left knee of the rats and were cut into small pieces. The meniscus pieces (100 mg) were digested with 3 mg/ml of collagenase and 4 mg/ml of dispase in 1 ml PBS at 37°C for 3 hours. Then the enzyme solution was removed by a centrifuge at 1500 g for 10 min, and the cell pellets were cultured in a tissue culture plate with 20% FBS-DMEM. The GFP cell numbers were examined under a fluorescent microscope (Nikon Eclipse E200, Japan).

### Histology analysis on wounded rat menisci

The meniscus tissues were collected from each group after irradiation, GFP-cell injection, and wound with different treatment. The meniscus tissues were fixed with 4% paraformaldehyde for 3 hours at room temperature and then decalcified with decalcifying solution (Sigma, St Louise, USA). After decalcification, the meniscus tissues were dehydrated through a series of graded ethanol baths and embedded with paraffin. Each tissue block was cut into 5 µm thick sections and stained with hematoxylin and eosin (H & E), and safranin O and fast green (S & F). The stained tissue sections were examined under microscope.

### Statistical analysis

All data were obtained from at least three replicates and presented as mean ± SD. The statistical analyses were performed with Excel 2007 version using student t-test or ANOVA and when P < 0.05, the two groups compared are considered to be significantly different.

## Results

### Preparation and characterization of HMGB1-KGN-AS bio-active scaffold

A novel bio-active scaffold for enhancing wounded meniscus healing has been developed using alginate gel to deliver HMGB1 and KGN to the wound area according to the scheme shown in **Fig. 1**. Sodium alginate was dissolved distilled water to make a 2% solution (AS), HMGB1 and KGN were added into AS to obtain a liquid mixture of HMGB1-KGN-AS (**Fig. 1A**). When the liquid mixture of HMGB1-KGN-AS was dropped into CaCl_2_ solution, a solid scaffold of HMGB1-KGN-AS was obtained by ion bond formation between alginates and Ca^2+^ (**Fig. 1B**). The control scaffold was made by adding the same volume of solvent (DMSO or/and water) instead of KGN or HMGB1 into AS.

**Fig. 1.**
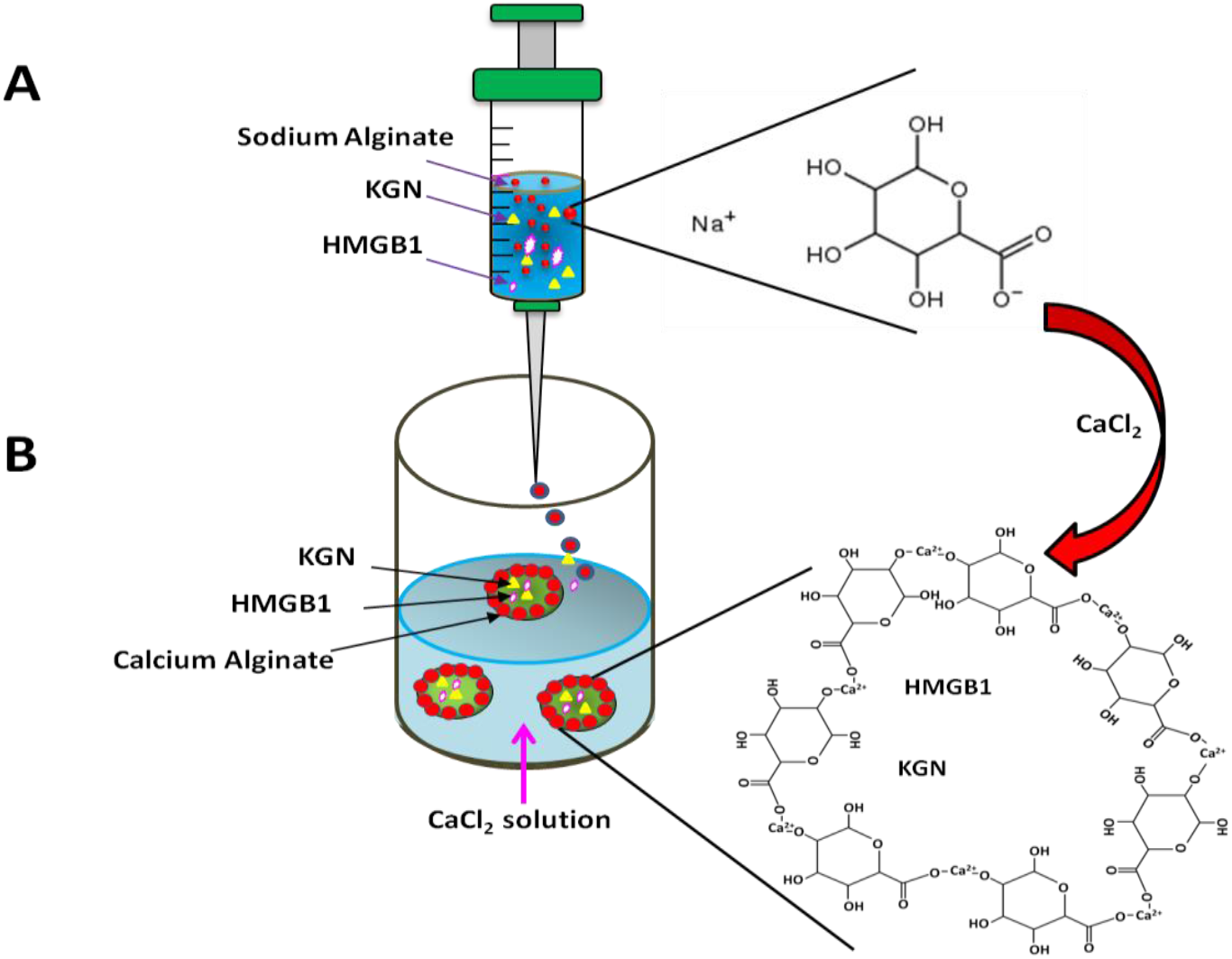
Scheme showed the preparation of bioactive scaffold. The bioactive scaffold was made by two steps. Step-1: KGN and HMGB1 were mixed with sodium alginate solution (**A**). Step-2: The KGN and HMGB1-containing sodium alginate solution was dropped into a calcium alginate solution to form a bio-activated scaffold (**B**). The structures showed that sodium alginate was a water soluble salt of alginic acid (**A**), however, when alginate was mixed with calcium ions, it is able to produce an insoluble polymer structure (**B**). Some drugs, such as HMGB1 and KGN were included in the inside of the calcium alginate (**B**).

Morphological analysis indicated that the control scaffold was a clear and empty bead (**Fig. 2A, 2C**), while KGN-containing scaffold was a solid bead with strong green fluorescence (**Fig. 2B, 2D**). HMGB1-containing scaffold was also a solid bead as evidenced by weak green fluorescence expression in the center area (**Fig. 2E**). Furthermore, the HMGB1 and KGN-containing bioactive scaffold was a solid bead with strongest green fluorescence (**Fig. 2F**).

**Fig. 2.**
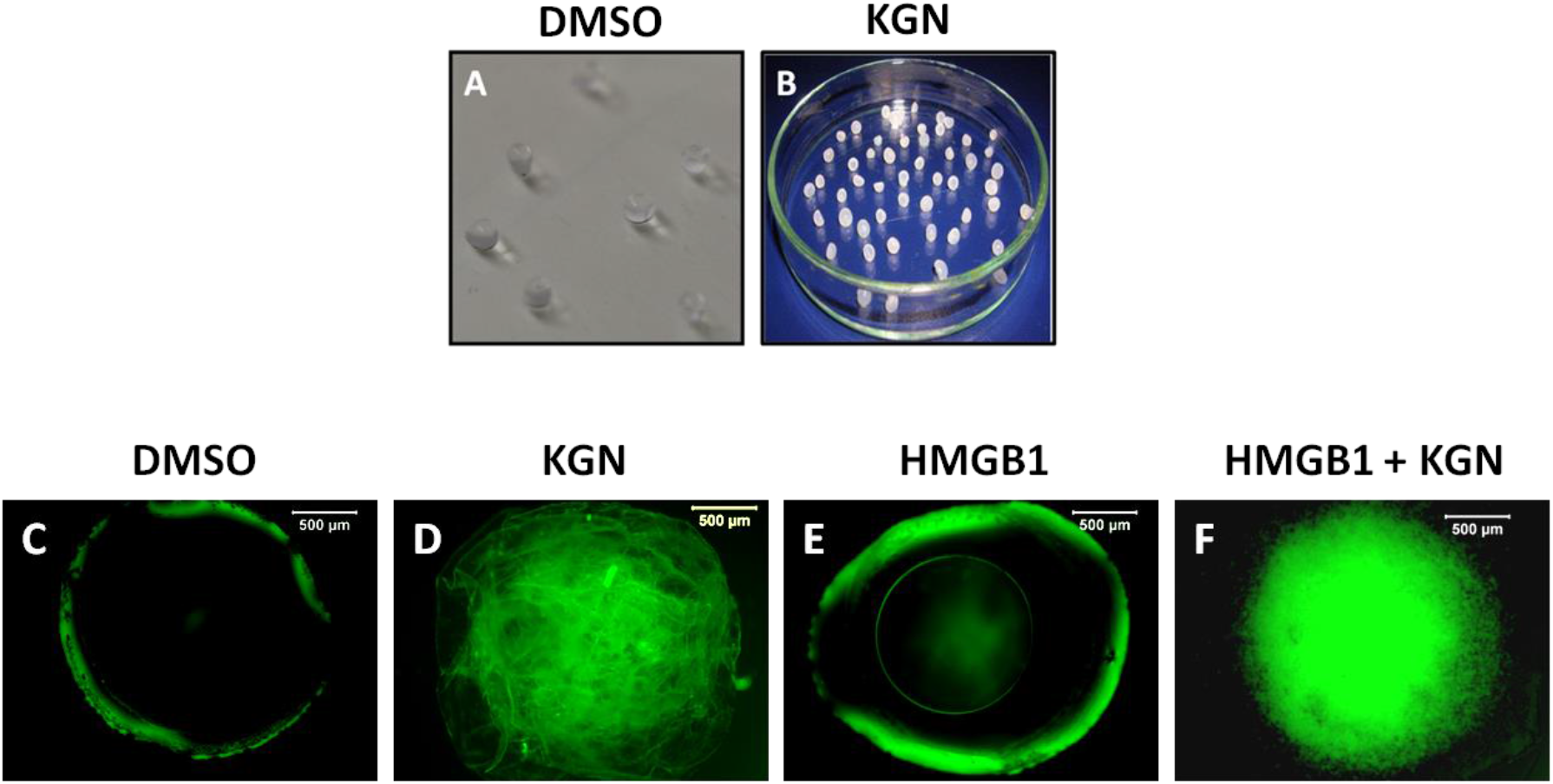
Gross and microscope view showed the morphology of bioactive scaffolds. **A, C:** DMSO-containing alginate scaffold was a clear and transparent bead (**DMSO**). **B, D**: The KGN-containing alginate scaffold was a white solid bead (**KGN**). **E:** The HMGB1-containing alginate scaffold (**HMGB1**). **F:** The HMGB1 and KGN-containing alginate scaffold (**HMGB1+KGN**). The KGN-containing beads showed solid beads with strong green fluorescence (**D, F**), while DMSO and HMGB1-containing alginate beads were empty beads with weak green fluorescence circle (**C, E**).

### Determination of HMGB1 and KGN released from HMGB1-KGN-AS bio-active scaffold

In vitro degradation testing results indicated that both HMGB1 and KGN can be released from the bio-activate scaffold made by HMGB1 and KGN-containing alginate gel in a time-dependent manner (**Fig. 3**). After 3 days, about 51% of HMGB1 (**Fig. 3A**) and 43% of KGN (**Fig. 3B**) have been released from HMGB1-KGN-AS scaffold, and kept high concentrations of HMGB1 and KGN for more than two weeks. At day-15, more than 90% of HMGB1 and KGN have been liberated from HMGB1-KGN-AS scaffold.

**Fig. 3.**
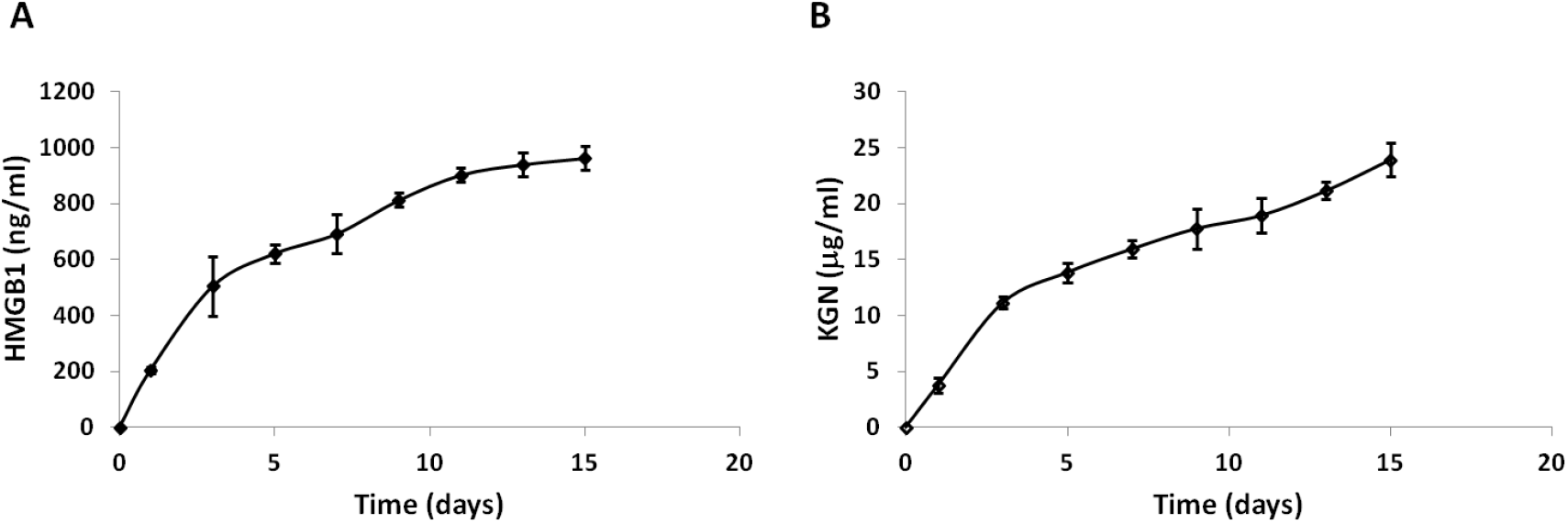
Drug release from bioactive scaffold in vitro tested in PBS. **A:** HMGB1 concentration released from HMGB1-KGN-containing alginate scaffold tested by ELISA kit. **B**: KGN concentration released from HMGB1-KGN-containing alginate scaffold in vitro tested in PBS by HPLC. Both HMGB1 and KGN were released from HMGB1-KGN-containing alginate scaffold in a time-dependent manner. After 3 days, more than 50% of HMGB1 and KGN have been released from the bio-active scaffold, and kept high concentrations of HMGB1 and KGN in the solution for two weeks. At day-15, more 90% of HMGB1 and KGN have been liberated from the HMGB1-KGN-AS scaffold.

### Effect of HMGB1 on cell migration

To determine the effect of HMGB1 on recruiting cell, a cell migration testing was investigated using a trans-well plate (**Fig. 4**).

**Fig. 4.**
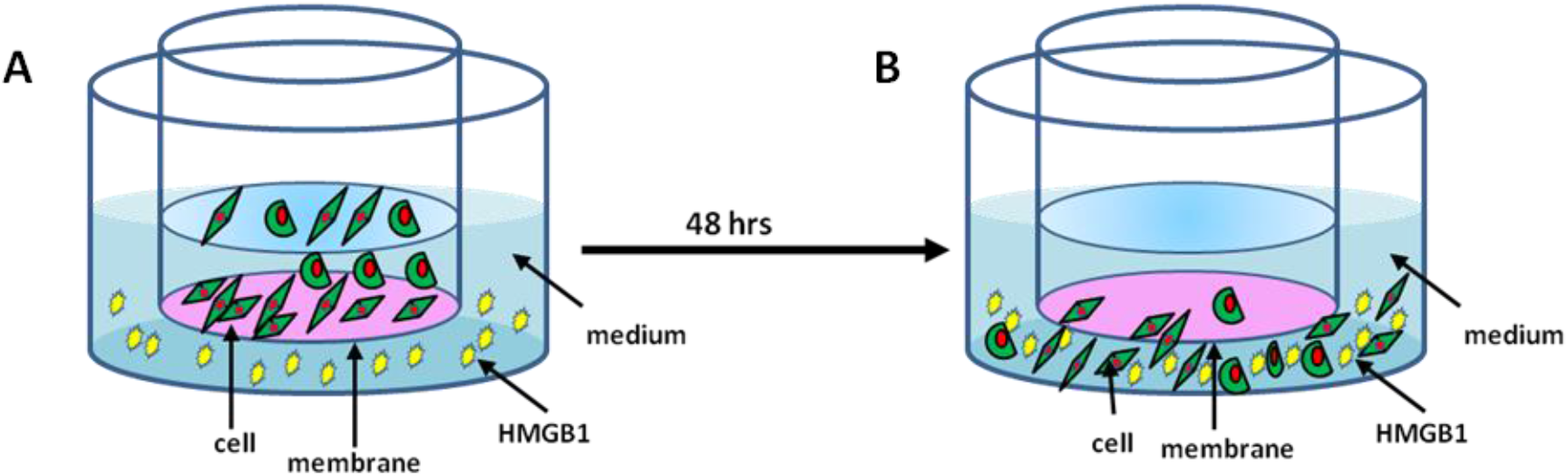
HMGB1 induced rat meniscus cell migration tested with a trans-well plate. **A:** Rat meniscus cells in serum-free medium at passage 2 were seeded in the upper layer of a trans-well plate insert with permeable membrane (5000 cells/well), and various concentrations of HMGB1 (0-100 ng/ml)-containing serum-free medium was placed below the cell permeable membrane. **B**: After 24 hours, the cells that have migrated through the membrane were stained with H33342 and counted under microscope.

The results showed that HMGB1 induced rat BMSC migration in a concentration-dependent manner (**Fig. 5A-E**). After 24 hours, very few cells migrated through the membrane into the serum-free medium without HMGB1 (**Fig. 5A, 5F**). However, more than 5 times of the cells have migrated through the membrane into the 100 ng/ml of HMGB1-containing serum-free medium (**Fig. 5D, 5F**) compared to the medium without HMGB1 (**Fig. 5A, 5F**).

**Fig. 5.**
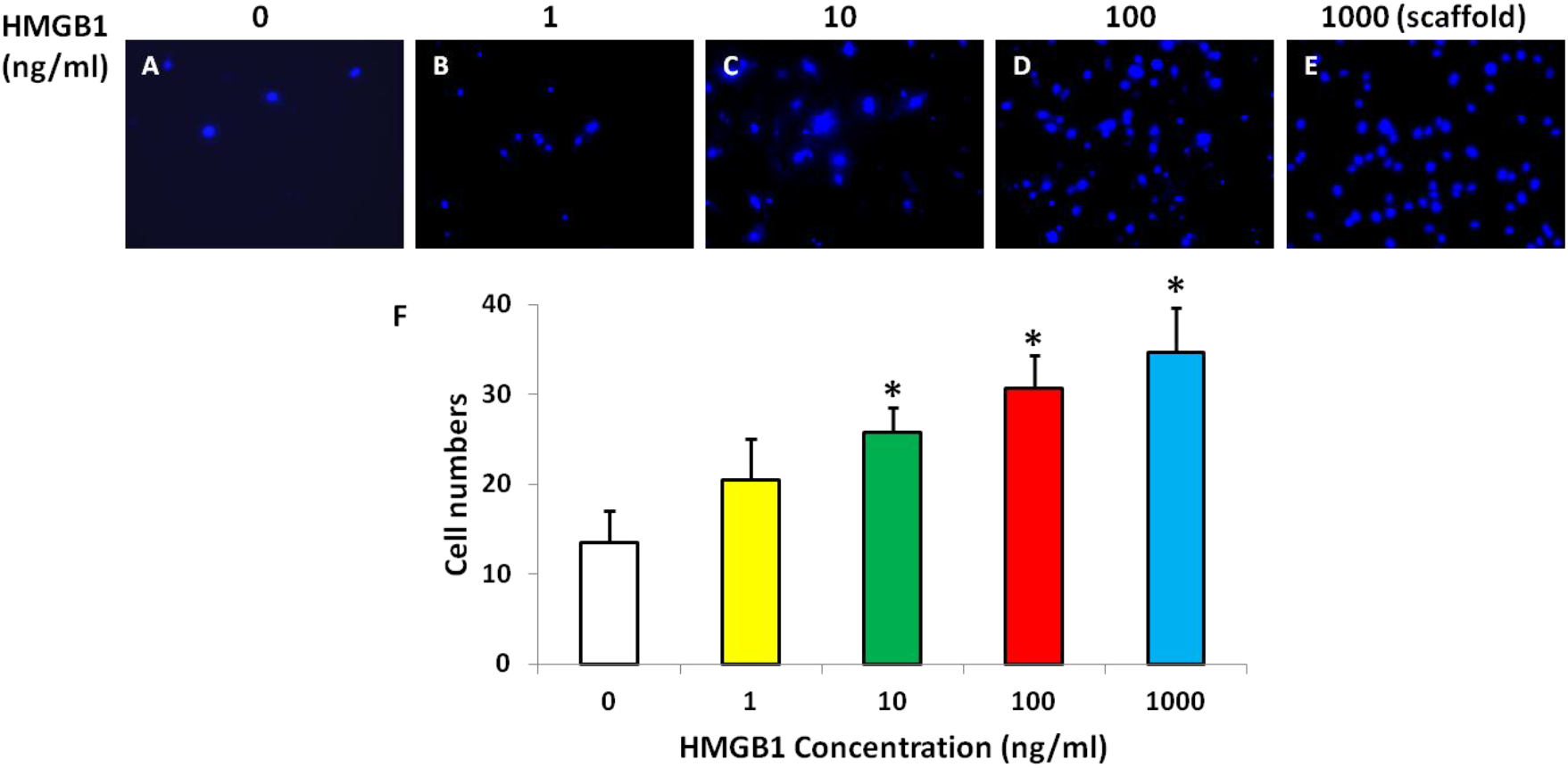
HMGB1 induced rat meniscus cell migration in a concentration-dependent manner. **A:** Cell migration was determined in serum-free medium with 0 ng/ml of HMGB1; **B:** Cell migration was determined in serum-free medium with 1 ng/ml of HMGB1; **C:** Cell migration was determined in serum-free medium with 10 ng/ml of HMGB1; **D:** Cell migration was determined in serum-free medium with 100 ng/ml of HMGB1; **E:** Cell migration was determined in 1 ml of serum-free medium with 1000 ng of HMGB1-containing AS scaffold; **F:** Semi-quantification of the cell migration was obtained from three trans-wells of each group. *p<0.05 compared to control (HMGB1 0 ng/ml).

The cell migration was inhibited by adding FPS-ZM1, an inhibitor of HMGB1 into HMGB1-containing medium (**Fig. 6**). The results indicated that FPS-ZM1 inhibited the recruiting effect of HMGB1 in a concentration-dependent manner as evidenced by the migrated cell numbers significantly decreased with the increased concentration of FPS-ZM1 in100 ng/ml of HMGB1-containg medium (**Fig. 6A-F**).

**Fig. 6.**
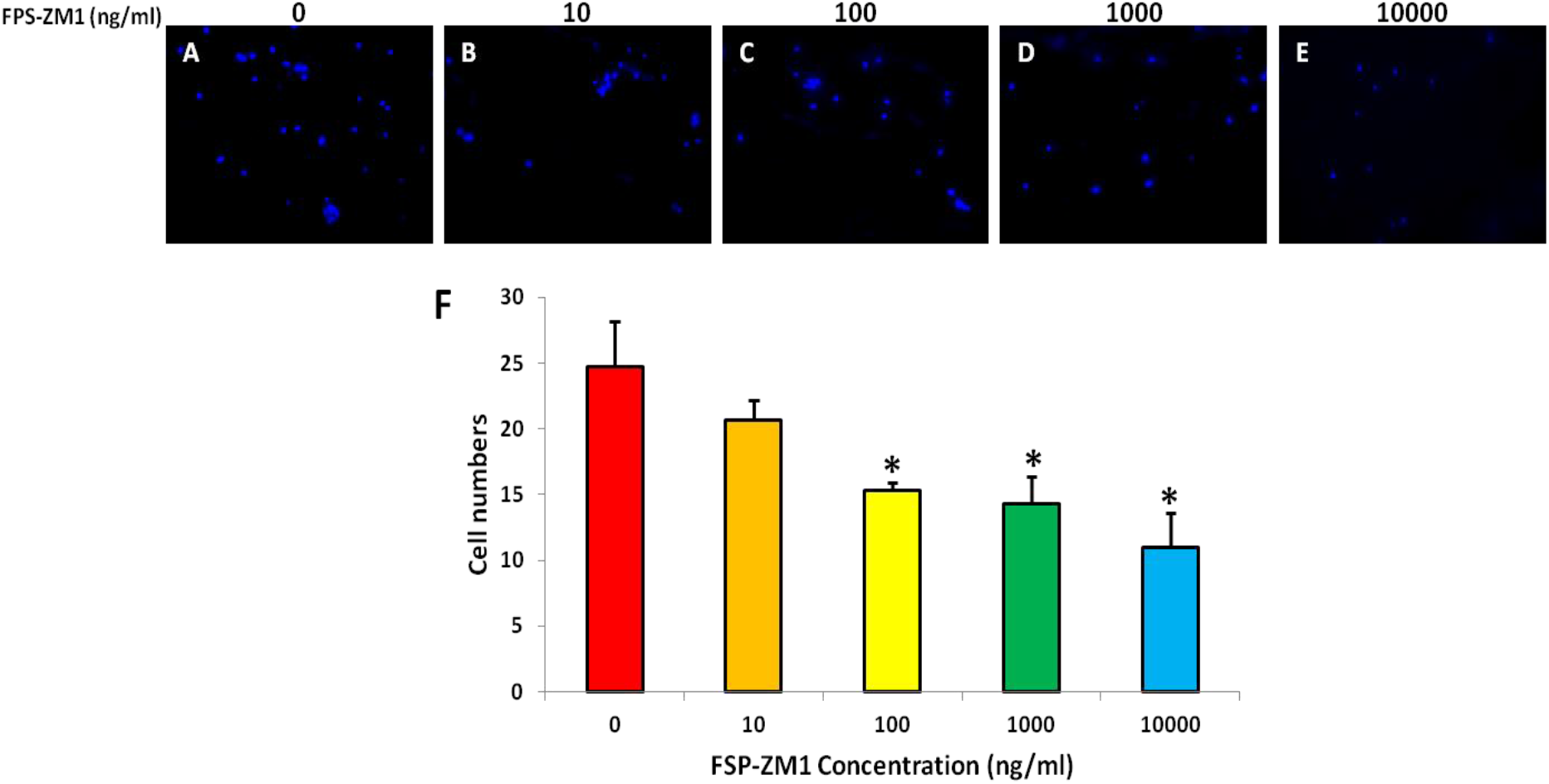
HMGB1 induced rat meniscus cell migration was inhibited by FPS-ZM1 in a concentration-dependent manner. Rat meniscus cells in serum-free medium at passage 2 were seeded in the upper layer of a cell culture insert with permeable membrane of trans-well plate (5000 cells/well), and various concentrations of FPS-ZM1 (0-10000 ng/ml)-containing serum-free medium with 100 ng/ml of HMGB1 was placed below the cell permeable membrane After 48 hours, the cells that have migrated through the membrane were stained with H33342 and counted under microscope. **A:** Cell migration was determined in serum-free medium with 100 ng/ml of HMGB1; **B:** Cell migration was determined in serum-free medium with 100 ng/ml of HMGB1 and 10 ng/ml of FPS-ZM1; **C:** Cell migration was determined in serum-free medium with 100 ng/ml of HMGB1 and 100 ng/ml FPS-ZM1; **D:** Cell migration was determined in serum-free medium with 100 ng/ml of HMGB1 and 1000 ng/ml of FPS-Zm1; **E:** Cell migration was determined in serum-free medium with 100 ng/ml of HMGB1 and 10000 ng/ml of FPS-Zm1; **F:** Semi-quantification of the cell migration was obtained from three trans-wells of each group. *p<0.05 compared to control (0 ng/ml of HMGB1).

Further experimental results confirmed these findings. The cell migration was also inhibited by adding another HMGB1’s inhibitor, AMD3100. Many cells have migrated through the membrane into HMGB1-containing medium without AMD3100 (**Fig. 7A, 7F**). The migrated cell numbers decreased significantly when AMD3100 was added into the HMGB1-containing medium, the more AMD3100, the less migrated cells (**Fig. 7**).

**Fig. 7.**
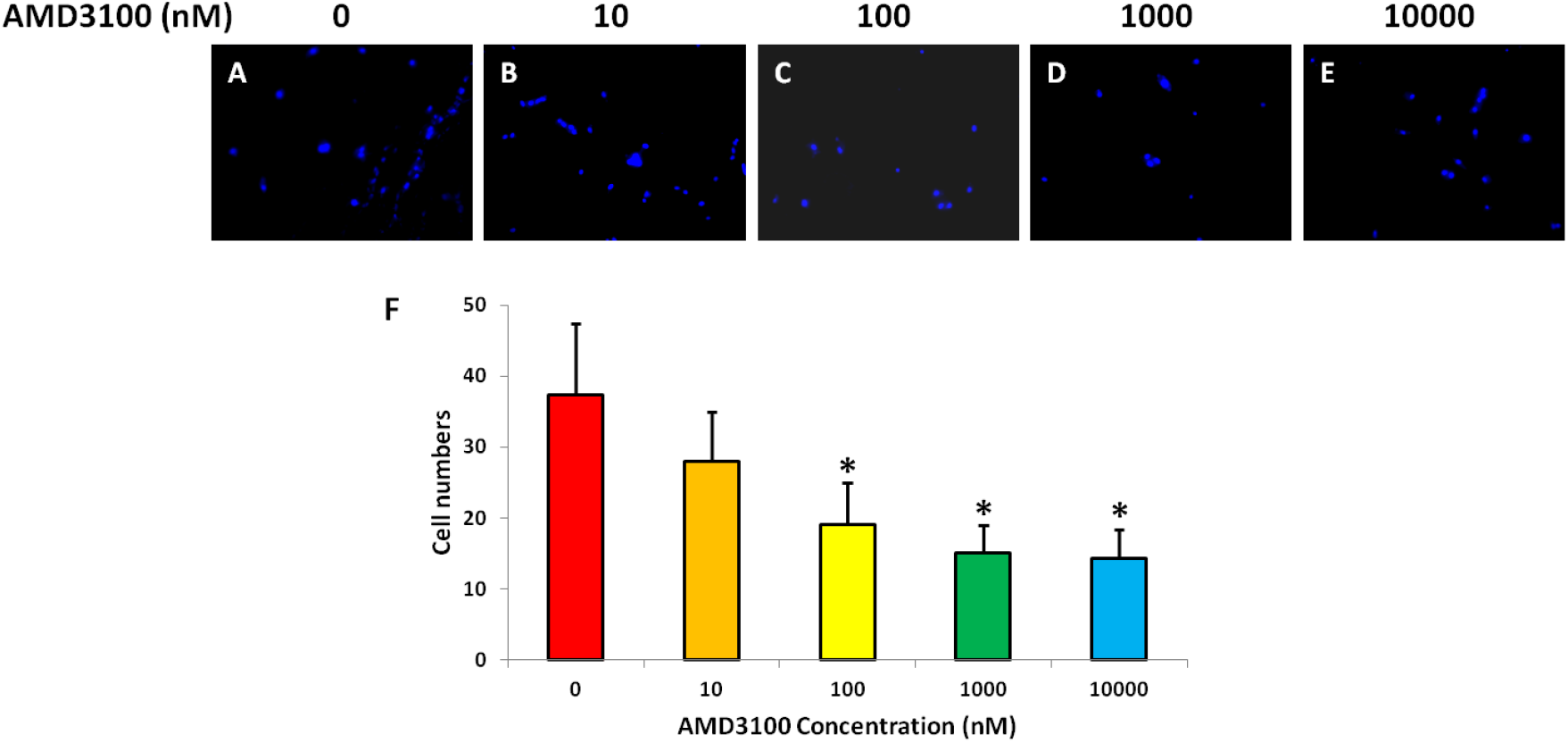
HMGB1 induced rat meniscus cell migration was inhibited by AMD3100 in a concentration-dependent manner. Rat meniscus cells in serum-free medium at passage 2 were seeded in the upper layer of a cell culture insert with permeable membrane of trans-well plate (5000 cells/well), and various concentrations of AMD3100 (0-10000 nM)-containing serum-free medium with 100 ng/ml of HMGB1 was placed below the cell permeable membrane After 48 hours, the cells that have migrated through the membrane were stained with H33342 and counted under microscope. **A:** Cell migration was determined in serum-free medium with 100 ng/ml of HMGB1; **B:** Cell migration was determined in serum-free medium with 100 ng/ml of HMGB1 and 10 nM AMD3100; **C:** Cell migration was determined in serum-free medium with 100 ng/ml of HMGB1 and 100 nM AMD3100; **D:** Cell migration was determined in serum-free medium with 100 ng/ml of HMGB1 and 1000 nM AMD3100; **E:** Cell migration was determined in serum-free medium with 100 ng/ml of HMGB1 and 10000 nM AMD3100; **F:** Semi-quantification of the cell migration was obtained from three trans-wells of each group. *p<0.05 compared to control (0 ng/ml of HMGB1).

### Effect of HMGB1 on cell proliferation

The HMGB1 effect on cell proliferation was also tested by BrdU staining. The results showed that frHMGB1 induced rat BMSC activation from G_0_ to G_Alert_ in a concentration-dependent manner (**Fig. 8**). In serum-free condition, more than 95% of the rat BMSCs were in G_0_ stage as evidenced very few cells were positively stained by BrdU (**Fig. 8A, 8K**). When frHMGB1 was added into serum-free medium, many rat BMSCs were activated from G_0_ to G_Alert_ stage as evidenced by cell numbers positively stained with BrdU (pink in **Fig. 8C-H**). Semi-quantification results indicated that more than 38% of the rat BMSCs in 1000 ng/ml of HMGB1-containing medium were positively stained BrdU (**Fig. 8K**).

**Fig. 8.**
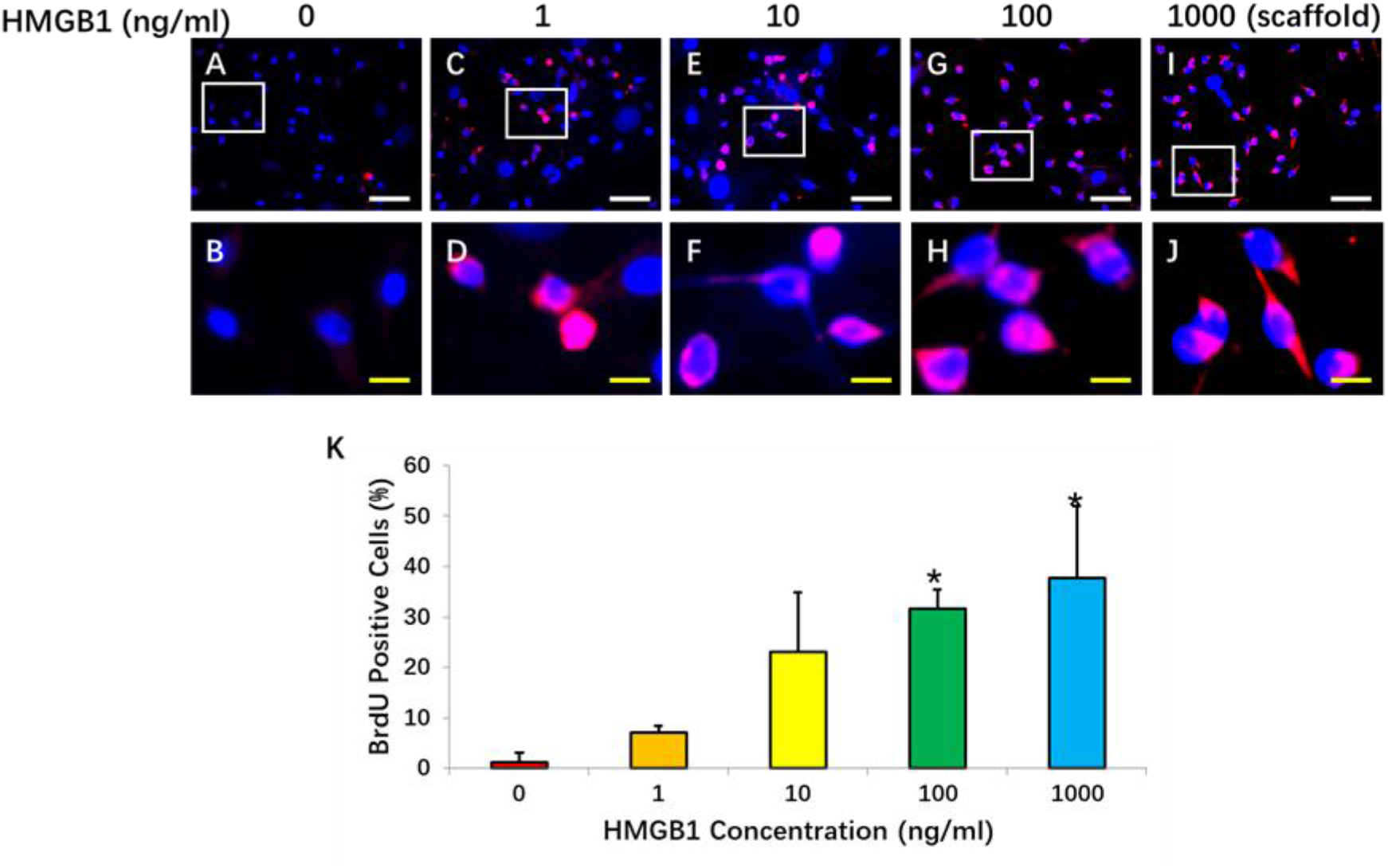
HMGB1 induced rat meniscus cell activation from G_0_ to G_Alert_ in a concentration-dependent manner. Rat meniscus cells at passage 2 were seeded in 12-well plate (20000 cells/well), and cultured with 20% of FBS-DMEM medium for 3 days. Then, the cells were cultured with serum-free medium for 24 hours, finally, various concentrations of HMGB1 (0-1000 ng/ml) with BrdU (10 mM) were added into the serum-free medium and the cells were cultured for another 24 hours. The cell proliferation was determined by anti-BrdU anti-body, and the **G_Alert_** cells were positively stained with BrdU indicating that BrdU has been incorporated into cells. Total cell numbers were stained with H33342 and the counted under microscope. **A, B:** The cells were cultured in 0 ng/ml of HMGB1; C, **D:** The cells were cultured in 1 ng/ml of HMGB1; **E, F:** The cells were cultured in 10 ng/ml of HMGB1; **G, H:** The cells were cultured in 100 ng/ml of HMGB1; **I, J:** The cells were cultured in 1000 ng HMGB1-containing AS scaffold in 1 ml serum free medium; **K:** Semi-quantification of the positively stained cells obtained from three wells of each group. The images of B, D, F, H, J were enlarged box areas in the images of A, C, E, G, I, respectively.*p<0.05 compared to control (HMGB1 0 ng/ml). White bars = 50 μm; Yellow bars = 10 μm.

### HMGB1-KGN-AS bioactive scaffold enhanced wounded meniscus healing

The effect of HMGB1-KGN-AS bioactive scaffold on promoting wounded rat meniscus healing was examined by a new *in vivo* model (**Fig. 9**). The native cells in the menisci of SD rats were eliminated using an irradiator (6 Gy) as shown in **Fig. 9A**. A wound was created by a biopsy punch (1 mm diameter) in the irradiated meniscus (**Fig. 9B**) and GFP-BMSCs were injected into tail vein of the irradiated rats (**Fig. 9C**). The wounds were treated with four different bioactive scaffolds (**Fig. 9D**), and the wounded meniscus healing results were examined 2 and 4 weeks post-operation by cellular analysis and histology studies.

**Fig. 9.**
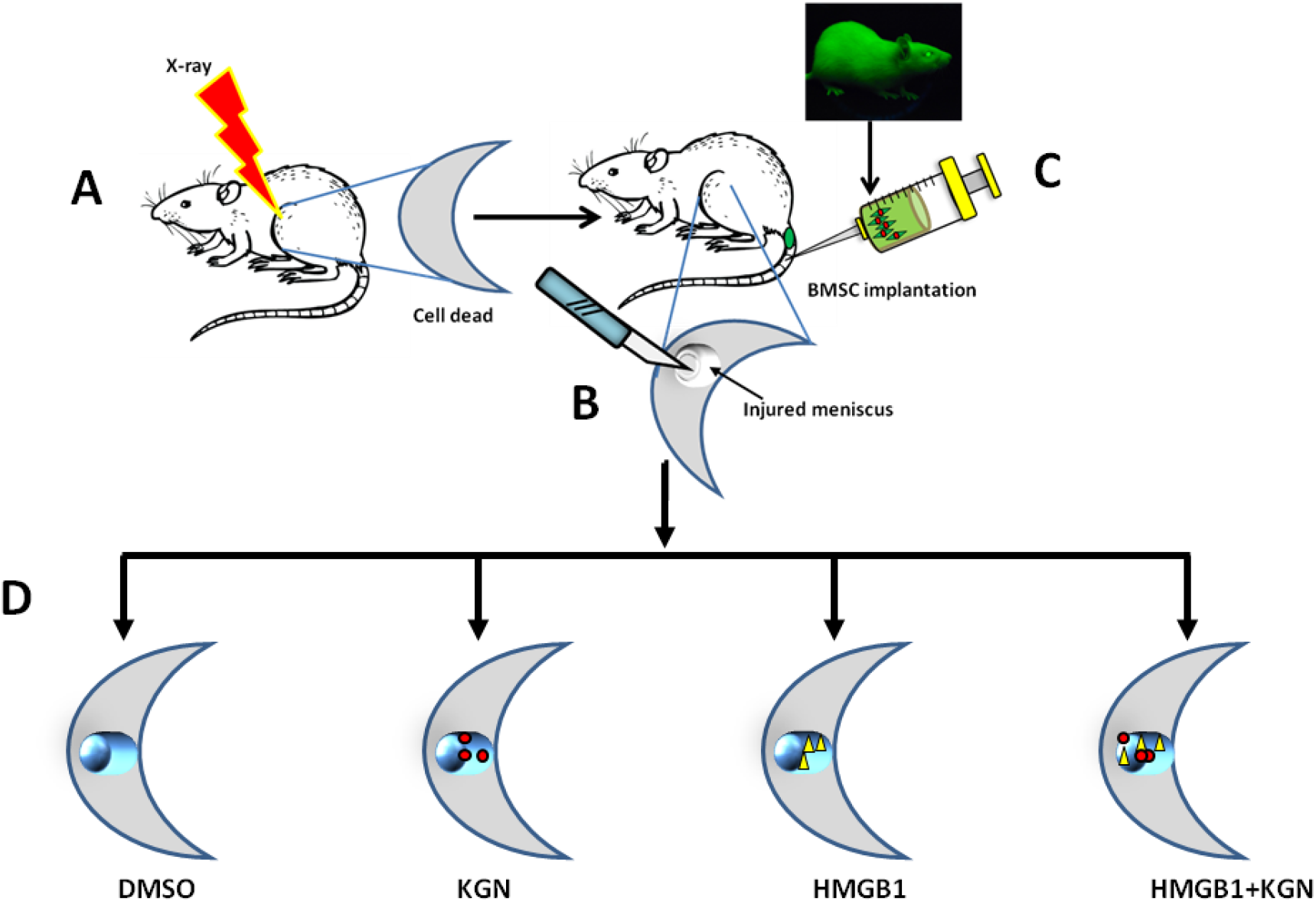
A new model showed that the bio-activated scaffold enhanced wounded rat meniscus healing by recruiting the cells to wound area by HMGB1 and differentiating the cells into chondrocytes by KGN. The rat meniscus cells were removed by irradiation, then green fluorescent protein-labeled rat bone marrow stem cells (BMSCs) were injected into tail vein of the rats. A wound (1 mm diameter) was created in the meniscus of the rats by a biopsy punch. The rats were divided onto four groups with 9 rats in each group. Group-1: The wound was treated with a DMSO-containing alginate bead (**DMSO**); Group-2: The wound was treated with a KGN-containing alginate bead (**KGN**); Group-3: The wound was treated with a HMGB1-containing alginate bead (**HMGB1**); Group-4: The wound was treated with a HMGB1 and KGN-containing alginate bead (**HMGB1+KGN**). The animals were sacrificed 2 and 4 weeks post-surgery. The effect of bio-activated scaffold on wounded meniscus healing was examined by cellular analysis and histology analysis.

The cellular analysis showed that the bioactive scaffold made by frHMGB1-KGN-AS can recruit cells to the wound area as evidenced by GFP cells found in the wounded menisci of SD rats (**Fig. 10**). The cells isolated from wounded rat menisci of all four groups expressed green fluorescence (**Fig. 10**), however, the cell numbers were significantly different. After two weeks healing, the GFP cell numbers in HMGB1-AS scaffold-treated meniscus (**Fig. 10C, 10M**) were 2 times as high as that in both DMSO-AS scaffold (**Fig. 10A, 10M**) and KGN-AS scaffold (**Fig. 10B, 10M**).

**Fig. 10.**
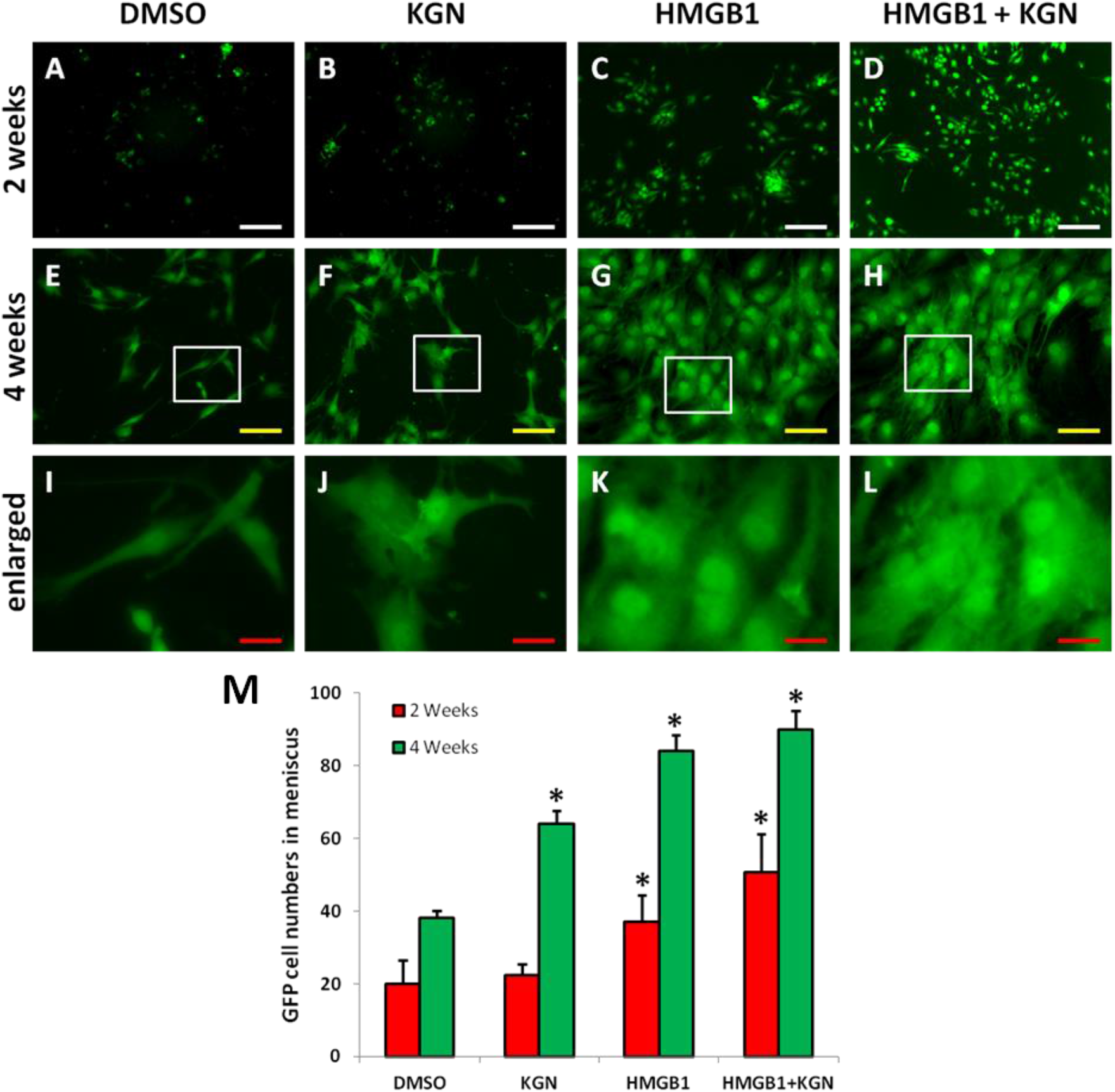
GFP cells isolated from wounded meniscus of the rats after various alginate beads treatments for 2 and 4 weeks. **A-D:**2 weeks; **E-L:**4 weeks; **A, E, I:** The wound was treated with a DMSO-containing alginate bead (**DMSO**); **B, F, J:** The wound was treated with a KGN-containing alginate bead (**KGN**); **C, G, K:** The wound was treated with a HMGB1-containing alginate bead (**HMGB1**); **D, H, L:** The wound was treated with a HMGB1 and KGN-containing alginate bead (**HMGB1+KGN**). **M:** Semi-quantification of GFP-cells calculated in each group. The meniscus was collected immediately after the rats were sacrificed, and the cells were isolated from the meniscus and cultured for 10 days according to our published protocol. The GFP-labeled cells in each group were examined under a fluorescent microscope. The results showed that HMGB1 recruited the GFP cells to migrate to the wound area. *p<0.05 compared to control (DMSC-AS scaffold treated wound). White bars: 200 μm; Yellow bars: 100 μm; Red bars: 25 μm.

Histology results further demonstrated that the bioactive scaffold made by HMGB1-KGN-AS gel not only recruited the cells to the wound area, but also induced the chondrogenic differentiation of BMSCs to enhanced wounded meniscus healing (**Fig. 11G, 11H**). The H & E staining results on tissue sections of wounded rat menisci showed that large unhealed wound areas were found in the wounded rat meniscus treated with DMSO-AS scaffold for 4 weeks (**Fig. 11A, 11B**). Although some cartilage-like tissues were found in the wounded rat meniscus treated with KGN-AS scaffold for 4 weeks (**Fig. 11C, 11D**), some unhealed wound areas were still presented in the wounded rat meniscus (**Fig. 11C, 11D**). High cell density was found in the wound areas of wounded rat menisci treated either with HMGB1-AS scaffold (**Fig. 11E, 11F**) or with HMGB1-KGN-AS scaffold (**Fig. 11G, 11H**), while more cartilage-like tissues were found in HMGB1-KGN-AS treated wounded rat meniscus (**Fig. 11G, 11H**).

**Fig. 11.**
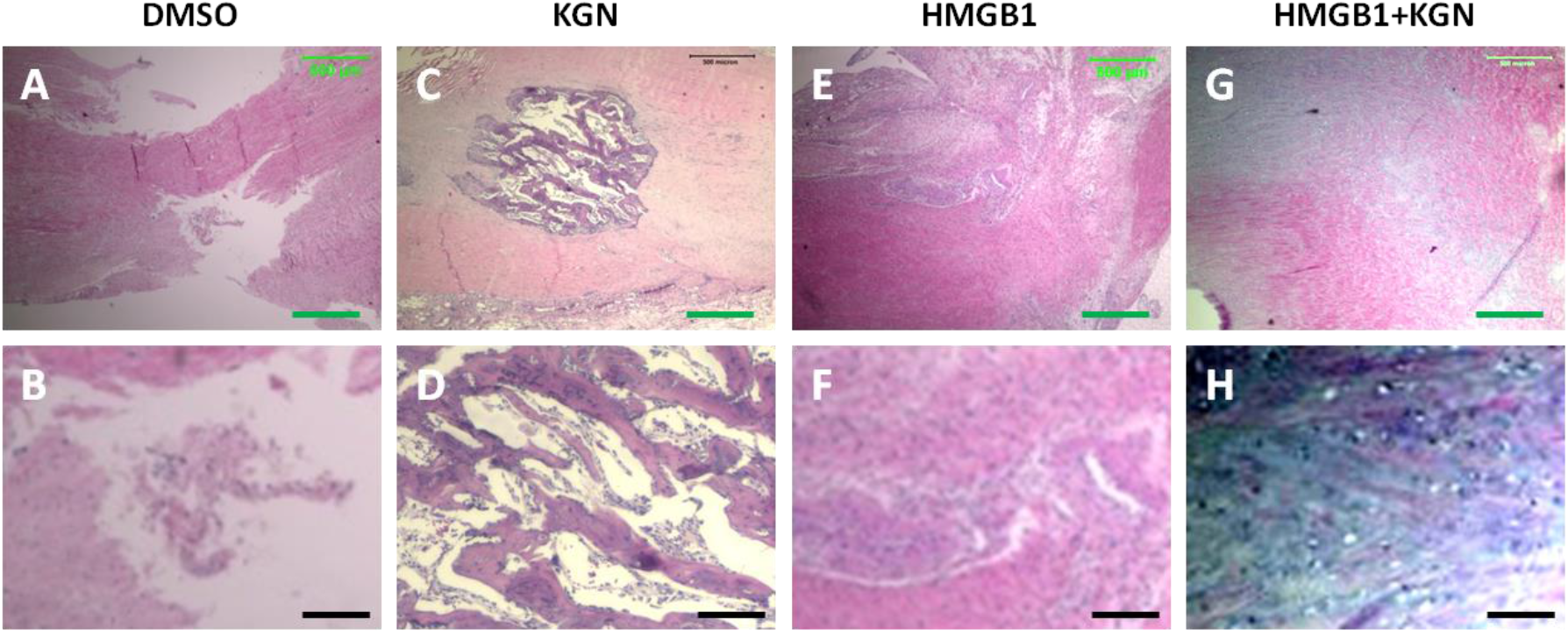
H & E staining on wounded meniscus of the rats after various alginate beads treatments for 4 weeks. **A, B:** The wound was treated with a DMSO-containing alginate bead (**DMSO**); **C, D:** The wound was treated with a KGN-containing bead (**KGN**); **E, F:** The wound was treated with a HMGB1-containing alginate bead (**HMGB1**); **G, H:** The wound was treated with a HMGB1 and KGN-containing alginate bead (**HMGB1+KGN**). Large unhealed wound area was found in DMSO-containing alginate bead treated wound (**A, B**). Some unhealed areas were still found in KGN-containing bead treated wound areas (**C, D**). The wound area was filled out with high density of the cells, but less cells were chondrocyte-like cells (**E, F**). The wound areas treated with HMGB1+KGN bead were filled with chondrocyte-like cells (**G, H**). Green bars: 500 μm; Black bars: 12.5 μm.

Further histology studies on safranin O and fast green staining confirmed H & E staining results (**Fig. 12**). Large unhealed areas were found in the wounded rat meniscus treated with DMSO-AS scaffold (**Fig. 12A, 12B**). Many chondrocyte-like cells in the wound areas of rat menisci treated either with KGN-AS scaffold (**Fig. 12C, 12D**) or with HMGB1-KGN-AS scaffold (**Fig. 12G, 12H**) were positively stained with safranin O, while the cell density in HMGB1-KGN-AS scaffold treated wounded area (**Fig. 12G, 12H**) was much higher than that of the wound area treated with KGN-AS scaffold (**Fig. 12C, 12D**). Some unhealed wound area was found in the wounded rat meniscus treated with KGN-AS scaffold (**Fig. 12C, 12D**). Although the wound treated with HMGB1-AS scaffold healed completely, very few cells were positively stained with safranin O (**Fig. 12E, 12F**).

**Fig. 12.**
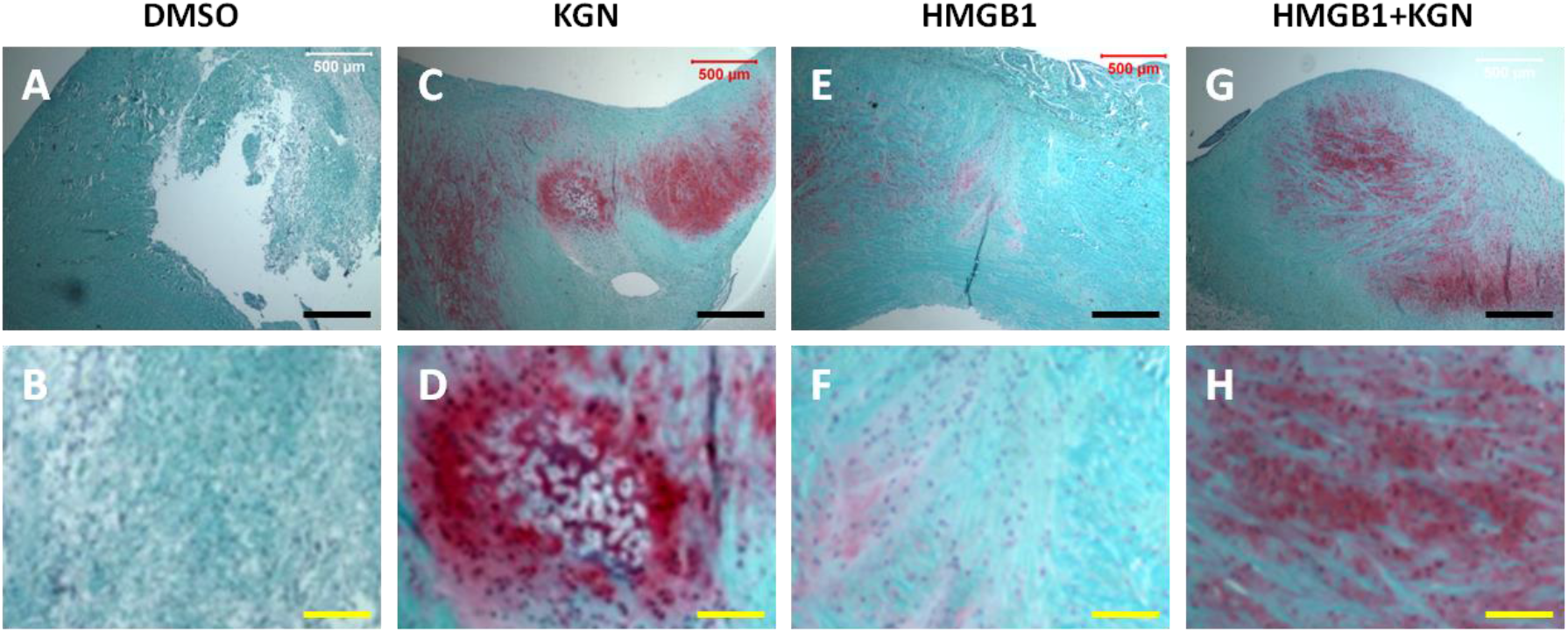
Safranin O and fast green staining on wounded meniscus of the rats after various alginate beads treatments for 4 weeks. **A, B:** The wound was treated with a DMSO-containing alginate bead (**DMSO**); **C, D:** The wound was treated with a KGN-containing bead (**KGN**); **E, F:** The wound was treated with a HMGB1-containing alginate bead (**HMGB1**); **G, H:** The wound was treated with a HMGB1 and KGN-containing alginate bead (**HMGB1+KGN**). Large unhealed wound area was found in DMSO-containing alginate bead treated wound (**A, B**). Although some unhealed areas were still found in KGN-containing bead treated wound areas, but the filled areas were positively stained by safranin O (**C, D**). The wound area was filled out with high density of the cells, but less cells were chondrocyte-like cells (**E, F**). The wound areas treated with HMGB1+KGN bead were filled with chondrocyte-like cells (**G, H**). Black bars: 500 μm; Yellow bars: 12.5 μm.

## Discussion

We have developed a novel bio-active scaffold for enhancing wounded meniscus healing by combination of frHMGB1 and KGN with alginate gel. The properties of the bio-active scaffold were also investigated *in vitro* and *in vivo*. We found that this HMGB1-KGN-AS bioactive scaffold released HMGB1 and KGN into wound area and kept high concentrations of HMGB1 and KGN in the system for more than two weeks. The results showed that the HMGB1-KGN-AS bioactive scaffold induced cell migration *in vitro* and promoted wounded meniscus healing *in vivo*.

Our results indicated that HMGB1 plays an important effect on recruiting cells to the wound area to enhance meniscus healing. The mechanism study demonstrated that HMGB1 recruited cells through RAGE signal pathway as evidenced by the inhibition effect of FPS-ZM1 on the HMGb1 induced cell migration. Our results demonstrated that the cell numbers in the bottom well were decreased significantly when FPS-ZM1 was added into HMGB1-containing medium. It has been reported that several receptors are implicated in HMGB1-mediated functions including receptor for advanced glycation end products (RAGE) [19]. Recent studies indicated that HMGB1 promotes intraoral palatal wound healing through RAGE-dependent mechanisms [20]. It is found that HMGB1 enhances recruitment of inflammatory cells to damaged tissues by forming a complex with CXCL12 and the signaling pathway is via CXCR4 [19]. It is known that FPS-ZM1 is a RAGE antagonist. Both in vitro and in vivo studies have shown that FSP-ZM1 blocked Aβ binding to the V domain of RAGE and inhibited Aβ40- and Aβ-42-induced cellular stress in RAGE-expressing cells and in the mouse brain [21]. It was recently demonstrated that HMGB1 is an enhancer of the activity of CXCL12 in stimulating the migration of mouse embryonic fibroblast [22].

Our similar results with AMD3100 further supported these findings. When we added AMD3100 into HMGB1-containing medium, the cell migration was significantly inhibited. It has been reported that AMD3100 is a CXCR4 antagonist and CXCL12-mediated engagement of CXCR4 works as an essential co-receptor signal for RAGE receptor-dependent HMGB1 migration response [23]. Our results revealed that the activity of HMGB1 is critical for cell migration.

The BrdU staining results showed that either HMGB1 along or HMGB1-containing bioactive scaffold accelerated the wounded rat meniscus healing by transitioning stem cells to G_Alert_ as evidenced by more BrdU-incorporating cells found in HMGB1-treated groups. BrdU has been used to label proliferating and migrating cells in various human and animal tissues and organs [24]. It has been found that BrdU competes with thymidine for incorporation into nuclear DNA during the S-phase of the cell cycle. Therefore, BrdU serves as a marker of DNA synthesis to ensure accuracy with regard to cell division [25].

This study also indicated that KGN is necessary for wounded meniscus healing due to it can induce the chondrogenic differentiation of stem cells. Safranin O staining results have shown that the wound treated with HMGB1-KGN-AS bioactive scaffold healed much faster than the other three groups and the wound area was filled with cartilage-like tissues. Although the wound treated with HMGB1-AS scaffold healed completely, there were a few chondrocyte-like cells found in the new formed tissues.

In this study, we used alginate to make bioactive scaffold. We found that alginate offered many advantages for wounded cartilage repair including its solubility in different ionic conditions and easy manufacturing process. Our results showed that sodium alginate is an injectable liquid, when sodium alginate solution was added into a calcium chloride solution, a solid scaffold was obtained. It has been reported that alginate is a biodegradable natural polymer which has a similar structure to the extracellular matrix (ECM) of chondrocytes [26]. Several studies have demonstrated that alginate provided an ideal environment to facilitate the spatial distribution of mesenchymal stem cells (MSCs), resulting in a structural organization that resembles the native *in vivo* cartilage microenvironment [17, 27]. Our results demonstrated that HMGB1-KGN-AS bioactive scaffold is beneficial for prospective applications in wounded meniscus repair.

